# Hallmarks of basidiomycete soft- and white-rot in wood-decay -omics data of *Armillaria*

**DOI:** 10.1101/2020.05.04.075879

**Authors:** Neha Sahu, Zsolt Merényi, Balázs Bálint, Brigitta Kiss, György Sipos, Rebecca Owens, László G. Nagy

## Abstract

The genus *Armillaria* spp. (Fungi, Basidiomycota) includes devastating pathogens of temperate forests and saprotrophs that decay wood. Pathogenic and saprotrophic *Armillaria* species can efficiently colonize and decay woody substrates, however, mechanisms of wood penetration and colonization are poorly known. We assayed the colonization and decay of autoclaved spruce roots using the conifer-specialists *Armillaria ostoyae* and *A. cepistipes* using transcriptomic and proteomic data. Transcript and protein levels were altered more extensively in the saprotrophic *A. cepistipes* than in the pathogenic *A. ostoyae* and in invasive mycelia of both species compared to their rhizomorphs. Diverse suites of carbohydrate-active enzyme genes (CAZymes), in particular pectinolytic ones and expansins, were upregulated in both species, whereas ligninolytic genes were mostly downregulated. Our gene expression data, together with previous comparative genomic and decay-chemistry analyses suggest that wood decay by *Armillaria* differs from that of typical white-rot fungi and shows features resembling soft rot. We propose that *Armillaria* species have modified the ancestral white-rot machinery so that it allows for selective ligninolysis based on environmental conditions and/or host types.

## Introduction

*Armillaria* spp. (Agaricales, Fungi) are among the most devastating fungal pathogens in woody ecosystems, including temperate forests, tree plantations, vineyards, and gardens [1–3] and are known to cause tremendous losses to the economy, health, and long-term productivity of forests [4–10]. The genus *Armillaria* is classified as white-rot fungi and comprises about 70 known species [11] including both pathogens and saprotrophs, making them suitable for studying mechanisms of pathogenicity and wood-decay systems in fungi [4, 8, 10–12].

*Armillaria* spp. can live both as plant necrotrophs and as saprotrophs [4, 10, 12, 13]. Based on their gene complement, they may decompose all components of wood, including the recalcitrant lignin [11–16] and other aromatic polyphenols along with cellulose, hemicellulose, and pectin [11, 12, 15, 16]. White rot fungi remove lignin from wood using high-redox potential oxidoreductases (e.g. class-II peroxidases [17–20]) and degrade complex polysaccharide polymers using diverse glycosyl hydrolase (GH), auxiliary activity (AA), carbohydrate esterase (CE) and polysaccharide lyase (PL) cocktails [20–23]. Decay-associated gene expression has been assayed in several species, mostly in the Polyporales (e.g. *Phanerochaete* spp. [24–28], *Ceriporiopsis subvermispora* [29], *Rigidoporus microsporus* [30, 31]*) and to a lesser extent in other clades (Heterobasidion* spp. [32–34], *Moniliopthora perniciosa* [35]). These studies highlighted wood species-specific responses, sequential activation of degradative enzymes, along with lifestyle-driven differences among species.

Previous comparative genomic studies on *Armillaria* have highlighted plant cell wall degrading enzyme repertoires reminiscent of white-rot fungi [11, 12, 15, 16], with a characteristic enrichment of pectinolytic genes [12, 16]. Accordingly, recent studies treated *Armillaria* spp. as white-rot based on the presence of lignocellulose degrading enzymes found in their genomes [11, 12, 16, 36, 37]. However, previous studies have also shown that *Armillaria* species primarily decay the cellulose, hemicellulose, and pectin components of the plant cell wall, and leave lignin unattacked during early stages of decay [38, 39]. Chemical and microscopic analyses of wood decay by *Armillaria* produced contradictory results. *A. mellea* was classified as a Group II white-rot fungi where celluloses and pentosans are decayed at early stages and lignin remains unaffected [39] resembling a possible brown-rot like approach. However, brown-rot is marked by an increased alkali-solubility in the wood aiding the fungi in the dissolution of plant cell wall carbohydrates, which was not found for *A. mellea* [39]. On the other hand, Schwarze et al reported a type-I soft-rot decay where the fungal hyphae grow through the secondary cell wall layer, producing characteristic cavities in the tracheids, axial and xylem ray parenchyma cells of Scots pine by *A. borealis, A. cepistipes, A. gallica, A*.*ostoyae* and *A. mellea* [38]. Soft-rot fungi by definition are now restricted to Ascomycota [14, 40–42], yet, there are many Agaricomycetes species that produce symptoms resembling soft rot or that did not fit the traditional white rot/brown rot dichotomy [36, 43, 44]. There are also reports suggesting a soft rot decay pattern in *Cylindrobasidium* spp [36] a close relative of *Armillaria* that also has a lower number of lignin-degrading enzymes as compared to typical white-rot decayers [12, 36, 45]. In order to place the *Armillaria* species into the ever-growing array of decay types, it is important to decode wood decay patterns in *Armillaria*.

In general, pathogenic basidiomycetes spread the infection by means of basidiospores. *Armillaria* spp. show another effective dispersal mechanism, through shoestring-like structures known as the rhizomorphs [4, 11]. Rhizomorphs are aggregations of hyphae exhibiting polarized apical growth, covered by a gelatinous sheath as their outermost layer. They serve as migratory or exploratory organs across larger distances beneath the soil [46–54]. It has also been speculated in previous studies that rhizomorphs might be responsible for foraging for nutrients [12, 46, 48] and for infecting plants via direct root contact [4, 9, 47, 55–57]. Rhizomorphs might also help *Armillaria* species become some of the largest and oldest organisms on Earth [58, 59]. At late stages of host colonization, rhizomorphs are often observed as thick, melanized cords on decayed and decorticated wood, however, their exact role in colonizing or degrading wood is poorly known.

We here employed a multi-omics approach to understanding wood-decay by *A. ostoyae* and *A. cepistipes*, both of which preferentially colonize conifers [13], the former as a pathogen while the latter as a saprotroph saprotroph with a mortality ratio not exceeding 5%, mostly in non-healthy trees [13, 60]. We allowed the two species to colonize sterilized Norway spruce roots (Fig 1A, B), then performed RNA-Seq and proteomics on invasive tissues (mycelium and rhizomorphs) and their non-invasive counterparts (*i*.*e*. mycelium and rhizomorphs grown in the absence of wood). Both species deployed a wide array of lignocellulose-degrading enzymes during root colonization, but the saprotroph *A. cepistipes* showed a stronger response to wood than did *A. ostoyae*. When compared with their non-invasive counterparts, invasive mycelia of both species harbored many more upregulated genes, including glycoside hydrolases, pectin-binding modules, carbohydrate-binding modules, hydrophobins, cytochrome P450s, and transcription factors than invasive rhizomorphs. We observed a decay pattern unusual for white-rot fungi, with weaker induction of lignin-targeting enzymes and upregulation of iron acquisition genes pointing towards a non-canonical white-rot strategy. Altogether our results shed light on soft rot and white rot wood-decay strategies in pathogenic and saprotrophic *Armillaria* species.

**Fig 1.**
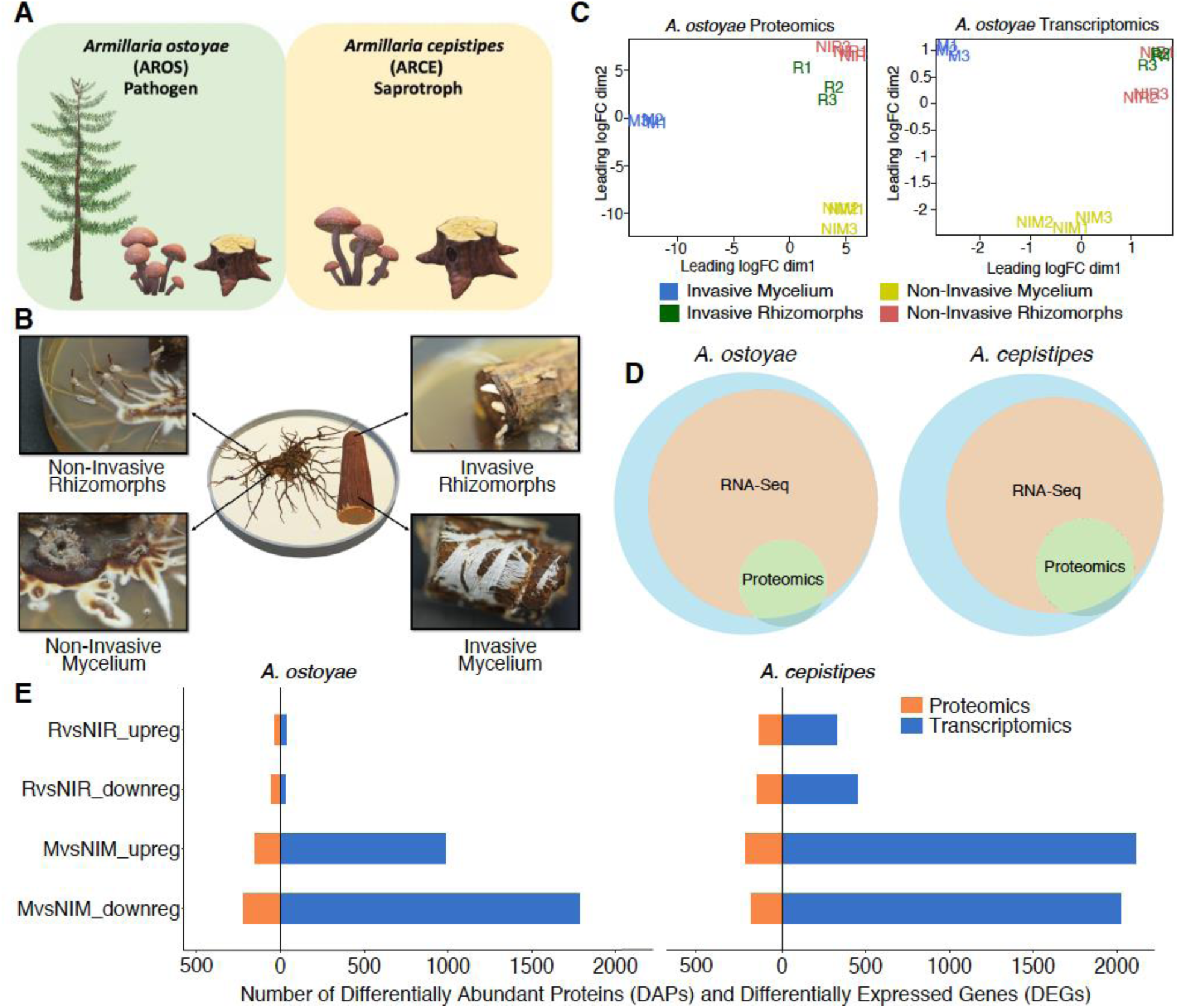
Overview of the experimental approach for root decay studies. A) Representation of *Armillaria ostoyae* (pathogenic) and *A. cepistipes* (saprotroph) used in this study, B) The four tissue types sampled for transcriptomics and proteomics analysis viz. invasive mycelium (growing beneath the outer layer of root), invasive rhizomorphs (emerging out of the roots), non-invasive mycelium and non-invasive rhizomorphs (growing in absence of root), C) Multidimensional scaling of three biological replicates from each of the tissue types in *A. ostoyae* for proteomics (left) and transcriptomics (right), D) Proportion of transcripts and proteins detected in the two -omics analysis. The blue circle represents the whole proteome, orange depicts the transcripts detected in the RNASeq, and green represents the proteins detected in the proteomics analyses. E) The number of differentially expressed genes (blue) and differentially abundant proteins (orange) detected in the two species.

## Results and Discussion

### Morphological observations and type of samples

Sterilized Norway spruce roots were introduced to one-week-old cultures of *A. ostoyae* and *A. cepistipes* and incubated at 25°C in the dark for 3-4 weeks until the roots were colonized. We observed an abundant growth of the mycelium in and under the bark layer (Fig 1B). Although previous studies suggested that colonization happens via direct rhizomorph contact and penetration [4, 9, 11, 12, 47, 55–57], we did not find evidence for the mechanical entry of rhizomorphs into the wood. Instead, upon coming in contact with the root, rhizomorphs switch to hyphal growth and spread further as mycelium (Fig S1-C). Rhizomorphs were not observed below the bark in *A. ostoyae* even after 8 weeks of incubation whereas *A. cepistipes* grew rhizomorph-like structures below the bark layer (Fig S1-B). Both species exited the root section as rhizomorphs emerging out of the piece of wood. These observations suggest that mycelium is the primary colonizing structure of wood, which is consistent with the higher surface/volume ratio of hyphae being better suited for nutrient acquisition as compared to rhizomorphs. Rhizomorphs probably emerge much later, possibly to transfer nutrients, as seen commonly under the bark of decayed logs [4].

### Overview of new -omics data

We analyzed four tissue types from both species in three biological replicates using transcriptomics and proteomics (Fig 1B). The mycelium and rhizomorphs collected from colonized roots are hereafter referred to as invasive mycelium (M) and invasive rhizomorphs (R), while those grown in the absence of roots are referred to as non-invasive mycelium (NIM) and non-invasive rhizomorphs (NIR), respectively. This yielded 12 samples for both *A. ostoyae* and *A. cepistipes* (with an additional sample type in the latter, see Fig S1-B). We prepared ribosomal RNA-depleted RNA libraries and sequenced them to a depth of 46.7-93.4 million paired-end reads on the Illumina NextSeq 500 platform. On average, 69% and ca. 38% of the reads mapped to the transcripts in *A. ostoyae* and *A. cepistipes*, respectively (Table S1). Concerning the low mapping percentages in *A. cepistipes*, we find that they were either caused by genomic DNA contamination or a poor annotation of the reference species (∼50% of unmapped reads mapped to intergenic regions, and not transcripts). Although such factors can dampen the signal of differential expression, we find that in our case this did not significantly compromise our analysis of differential gene expression (see the clear separation of samples in the MDS and the high number of DEGs, Fig 1C, E).

Label-free comparative proteomics provided relative abundance of data across the different sample types. A total of 37,879 and 36,087 peptides were identified from *A. ostoyae* and *A. cepistipes*, respectively, which were subsequently rolled up into protein groups, with median protein sequence coverage ranging from 30.1 – 34%.

Multidimensional scaling (MDS) plots show a strong clustering of the biological replicates in both transcriptomic and proteomic data in both species. The MDS plots portray a clear separation of the invasive and non-invasive mycelium samples whereas the invasive rhizomorphs and non-invasive rhizomorphs showed higher similarity to each other (Fig 1C for *A. ostoyae*, Fig S1 for *A. cepistipes*), suggesting that the larger difference exists between invasive and non-invasive mycelium. For *A. ostoyae*, RNA-Seq detected and quantified 17,198 transcripts, while the label-free proteomics analysis detected 3,177 proteins. In *A. cepistipes* we obtained data for 15,861 transcripts and 3,232 proteins (Fig 1D). Overlap between the two - omics datasets was substantial about 99.8% of the detected proteins to be also present in the RNA-Seq data (Fig 1D). At the same time,there was a limited correlation between the fold-change values acquired for transcripts and proteins (Fig S2), which is not unusual among proteomic and transcriptomic datasets.

Expression analyses were carried out to identify differentially expressed genes (DEGs). In *A. ostoyae* we found 987 and 35 upregulated genes (log_2_FC>1, p-value<0.05) in invasive vs. non-invasive mycelium (M*vs*NIM) and in invasive vs. non-invasive rhizomorphs (R*vs*NIR) (5.74% and 0.2% of transcriptome), respectively. Considerably more, 2,108 and 327 upregulated genes were found to be differentially expressed in M*vs*NIM and R*vs*NIR in *A. cepistipes*, respectively (13.29% and 2.06% of transcriptome; Table S2).

The number of differentially abundant proteins (DAPs) in the proteomics analysis was lower: we detected 279 and 108 proteins with increased abundance in MvsNIM and RvsNIR in *A. ostoyae* (8.78 and 3.40 % of detected proteome), and 439 and 282 proteins with increased abundance in MvsNIM and RvsNIR, respectively in *A. cepistipes* (13.58 and 8.73 % of detected proteome; Table S3).

We found considerably more DEGs/DAPs in the mycelium than in rhizomorphs (Fig 1E) in both species, which probably indicates that the mycelium is more actively involved in the colonization of woody tissues than are rhizomorphs. This is consistent with the morphological observations and an invasion of the roots primarily by individual hyphae. A more surprising observation is that the saprotrophic *A. cepistipes* shows a higher number of DEGs/DAPs than the pathogenic *A. ostoyae* (Fig 1E). To confirm that this is not a result of higher baseline expression of some genes in *A. ostoyae*, we compared the distribution of raw expression values of co-orthologs in the M and NIM samples in both species (Fig S3). This showed that baseline expression of genes in non-invasive mycelia of *A. ostoyae* was not higher than that in *A. cepistipes*, indicating that the higher number of DEGs in *A. cepistipes* is indeed the result of the stronger reaction of this species to wood. We speculate that this is because saprotrophs, to gain a competitive advantage over other microbes, have to colonize/degrade wood faster than necrotrophic pathogens, such as *A. ostoyae*, which can both feed on living parts of the tree and, upon killing the host, can be the first colonizers of the wood [12, 61].

### Gene ontology (GO) analyses

In the transcriptomics data for *A. ostoyae*, we found 25 and 11 GO terms significantly enriched among the genes upregulated in M*vs*NIM and R*vs*NIR, respectively (Fig 2, left). For *A. cepistipes* we found 57 and 29 terms enriched in M*vs*NIM and R*vs*NIR, respectively (Fig S4, left). In both species, genes upregulated in invasive mycelia were enriched (p<0.05, Fisher’s exact test) for terms related to oxidation-reduction, lipid metabolism, transmembrane transport, iron ion and heme-binding processes along with the terms ‘fungal type cell wall’ and ‘structural constituent of cell wall’. We find that the enrichment of iron-binding related terms was driven by the upregulation of members of the high-affinity iron permease complex (2 out of 3 genes and, 1 out of 2 genes upregulated in *A. ostoyae* and *A. cepistipes*, respectively, see below). Consistent with the lower number of DEGs in invasive vs non-invasive rhizomorphs, we observed fewer enriched GO terms (see Table S4).

**Fig 2.**
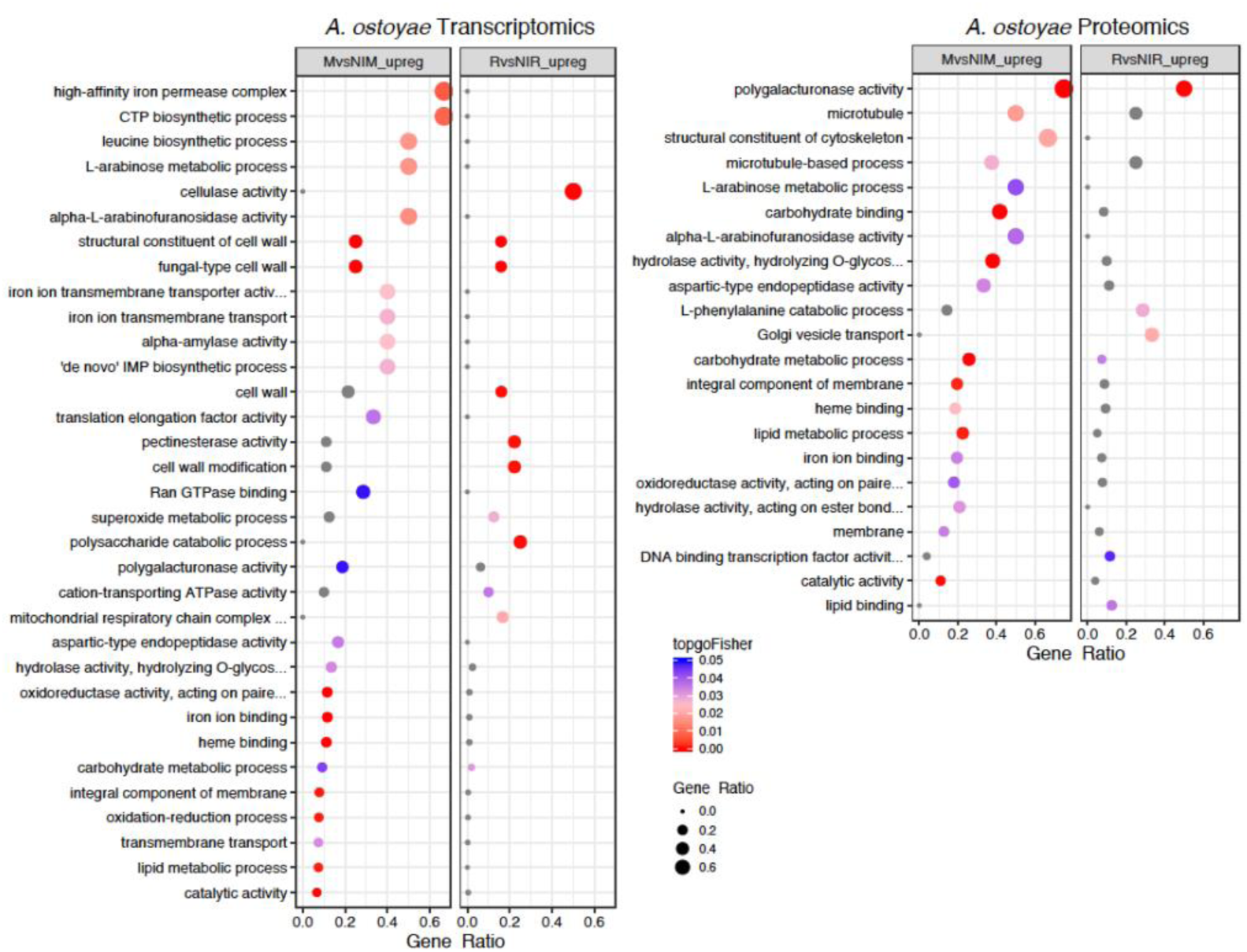
Enriched GO terms in M*vs*NIM and R*vs*NIR of *A. ostoyae* for transcriptomics (left) and proteomics (right). The ratio of number of a particular GO term in a specific comparison (mycelium vs non-invasive mycelium or in rhizomorphs vs non-invasive rhizomorphs) to the total number of that GO term for a species was used to plot gene ratios for enriched GO terms (p<0.05, Fisher’s exact test). The size of the dot is directly proportional to gene ratio, and the color of the dots corresponds to p-values. Grey dots represent GO terms, enriched in only one of the comparisons *i*.*e* either mycelium vs non-invasive mycelium or rhizomorphs vs non-invasive rhizomorphs. Enriched GO terms in M*vs*NIM and R*vs*NIR of *A. cepistipes* can be found in Fig S2.

We observed a similar pattern in the proteomic data, there were more enriched GO terms in *A. cepistipes* than *A. ostoyae*. In *A. ostoyae*, we found 18 and 12 significantly enriched terms in mycelium vs non-invasive mycelium and rhizomorphs vs non-invasive rhizomorphs, respectively (Fig 2 right), whereas, in *A. cepistipes*, there were 35 significantly enriched terms in mycelium vs non-invasive mycelium and 19 in rhizomorphs vs non-invasive rhizomorphs (Fig S4, right). Terms enriched in mycelium vs non-invasive mycelium included pectinesterase activity, polygalacturonase activity, cellulose-binding, carbohydrate-binding, hydrolase activity, together with cell wall modification, cell wall-related terms, sugar metabolic processes.

### Global transcriptome and proteome similarity

We measured global similarity among transcriptomes and proteomes within and across species based on Pearson correlation. In general, we observed a better correlation among transcriptomes than among proteomes (Fig 3).

**Fig 3.**
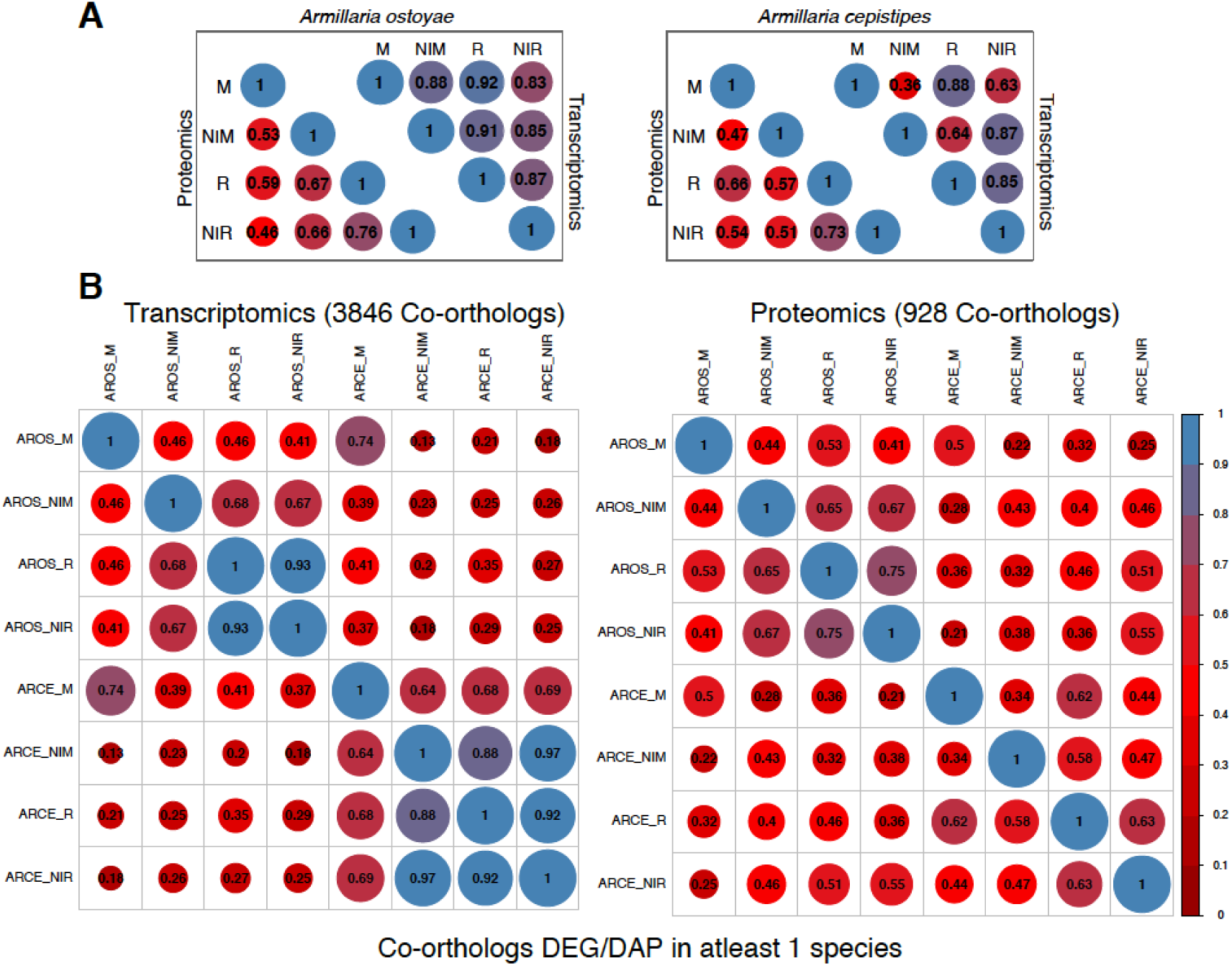
Global transcriptome/proteome similarity between *A. ostoyae* and *A. cepistipes*. A) Correlation between the 4 tissue types for proteomics and transcriptomics data in *A. ostoyae* (left) and *A. cepistipes* (right). B) Correlation between co-orthologs which were significantly differentially expressed/abundant in at least one of the species (for all co-orthologs, see Fig S3), showing correlation between samples across the two species. Blue represents higher correlation and red represents lower. Size of the circle is directly proportional to higher correlation. Pairwise mean Pearson correlation coefficients are given as numbers in the circles.

Within species, we observed limited differences among transcriptomes, with the highest global transcriptome similarity values observed between invasive tissue types (Fig 3A). For example, in *A. ostoyae*, the two most similar sample types were invasive mycelium and invasive rhizomorphs (mean Pearson: 0.92), slightly higher than other combinations of samples (0.83 - 0.88). A similar, but a stronger pattern is observable in *A. cepistipes* (Fig 3A). This pattern suggests that contact with wood elicits similar expression changes irrespective of the tissue type. In support of this, we could identify 4 and 127 genes upregulated in the invasive mycelium and invasive rhizomorphs of *A. ostoyae* and *A. cepistipes*, respectively. Many of these genes were annotated as hydrophobins, cytochrome P450s, galactose-binding domain-like proteins, and a number of CAZymes (Table S2).

The among-species similarity between sampled tissues was assessed based on 11,630 co-orthologous genes in *A. ostoyae* and *A. cepistipes*, identified by OrthoFinder [62], out of which transcriptomic and proteomic data cover 10,675 and 2,404, co-orthologs, respectively. A surprisingly high correlation was found between the invasive mycelia of *A. ostoyae* and *A. cepistipes* (Fig S5), whereas the correlation was comparatively lower in all other combinations. This observation was similar for both transcriptomic and proteomic data and was even more pronounced when we considered only genes/proteins that were DEG or DAP in at least one of the species (Fig 3B). We interpret the correlated gene expression in *A. ostoyae* and *A. cepistipes* as an indication of a shared response of invasive mycelia to the presence of spruce roots.

### Shared transcriptomic response of mycelia to wood

To understand what comprises the observed similarity in wood decay, we focused on co-orthologous gene/protein pairs up- or downregulated in invasive mycelia of both species. We found 779 co-orthologs having similar differential expression in the invasive mycelium. Of these 372 and 407 were significantly up- and downregulated, respectively (Fig 4B, Table S5). For the 372 upregulated co-orthologs, we observed overall higher fold changes and expression levels in *A. cepistipes* than in *A. ostoyae* (Fig 4B), again, underscoring a stronger response of *A. cepistipes* to wood.

**Fig 4.**
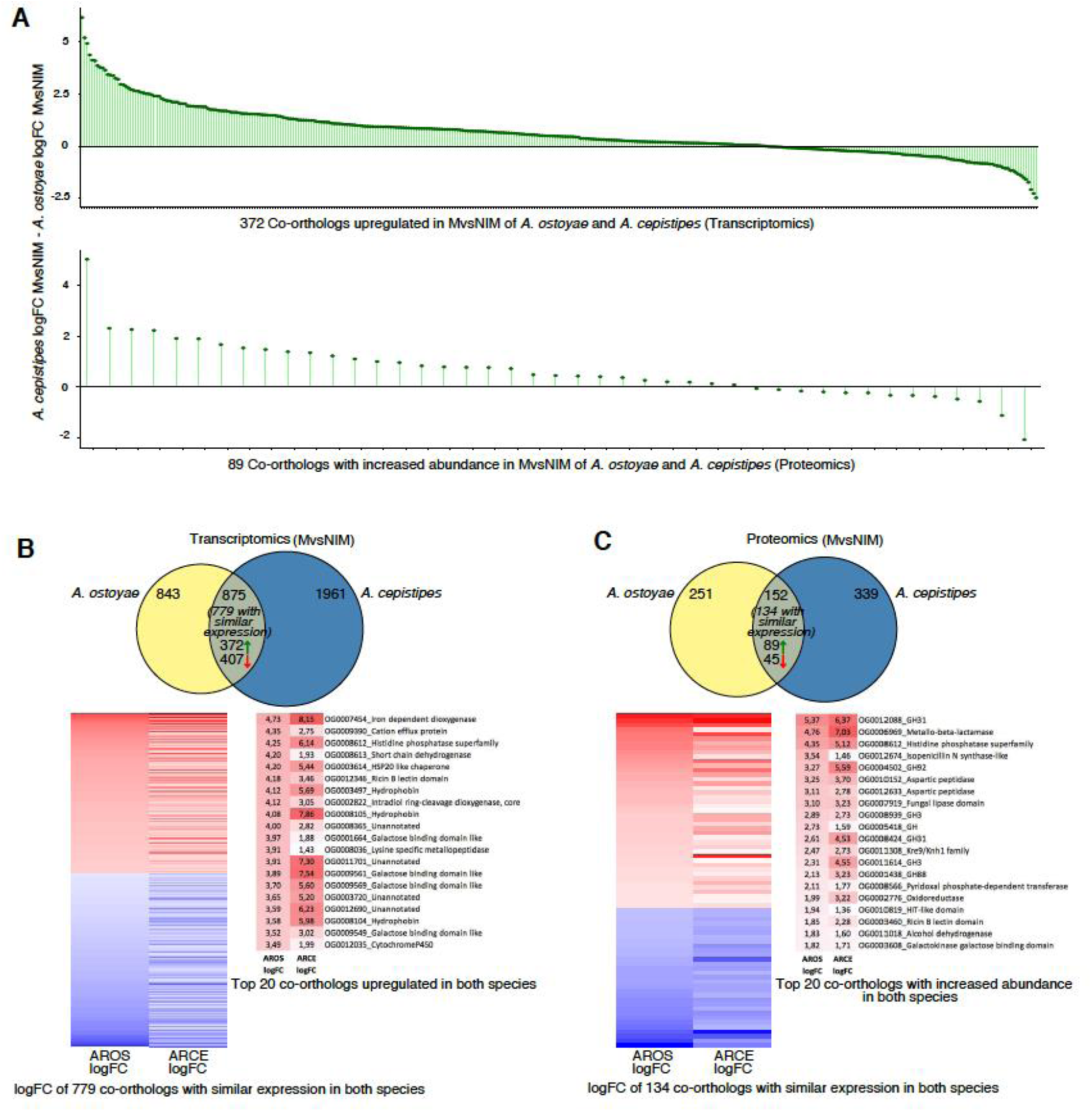
Response towards spruce roots by the mycelia (MvsNIM) of *A. ostoyae* and *A. cepistipes*. A) logFC differences of co-orthologs DEG (up) and DAP (down) in both species, For proteomics, only genes for which a fold change could be calculated are shown (43 out of 89 orthologs). B) Venn diagram showing species-specific and common DEGs in *A. ostoyae* and *A. cepistipes*. Heatmap below showing 779 co-orthologs with similar expression patterns (upregulated/downregulated in both species) out of the 875 common DEGs. Towards right are the top 20 upregulated proteins (red) in the two species. C) Venn diagram showing species-specific and common DAPs in *A. ostoyae* and *A. cepistipes*. Heatmap below showing 134 co-orthologs with similar abundance (increased/decreased in both species) out of the 152 common DAPs Towards right are the top 20 proteins with increased abundance in the two species.

Among the most upregulated co-orthologs in the transcriptomic analyses, we found oxoglutarate/iron-dependent dioxygenases, proteins of the galactose-binding-like domain superfamily (including CBM67, see below), ricin-B lectins, hydrophobins, intradiol-ring cleavage dioxygenases, GMC oxidoreductases, cytochrome p450-s, as well as 10 conserved transcription factors and several unannotated genes (Fig 4B, Table S5). The most highly induced genes in both species were oxoglutarate/iron-dependent dioxygenases. These were reported to be responsible for the oxidation of organic substrates, mycotoxin production, and secondary metabolite biosynthesis [63–65] and were also found to be upregulated in both white-rot and brown-rot wood decay studies [66–68]. We found 46 oxoglutarate/iron-dependent dioxygenase genes in both species, of which 5 and 9 were upregulated in the invasive mycelium of *A. ostoyae* and *A. cepistipes*, respectively (but not in invasive rhizomorphs). In proteomics, we found 1 and 4 genes in the invasive mycelium and 1 and 2 genes in the invasive rhizomorphs with increased abundance in *A. ostoyae* and *A. cepistipes*, respectively. The 2-oxoglutarate dioxygenase superfamily is widespread across microorganisms, fungi, plants, and mammals as well [65, 69–71], however, their versatile nature makes them difficult to interpret in terms of exact biological relevance in wood decay mechanisms.

In the proteomics data, we found 89 co-orthologs with increased and 45 with decreased abundance in both species (Fig 4C, Table S5), of which, the ones with the highest abundance in both species included GH31, GH3, GH88, GH92 CAZyme families as well as aspartic peptidases, fungal lipases and Kre9/Knh1 fungal cell wall-related proteins. Some of these proteins were only detectable in the invasive mycelium and not in non-invasive mycelia, including several CAZymes such as pectin lyases, GH28, carbohydrate-binding modules (CBMs), PL8, GH3, GH35, galactosidases, carboxylesterases and several other gene families like GMC oxidoreductases, various transporters, cytochrome P450s.

#### Characteristic PCWDE expression in invasive mycelia

A diverse array of plant cell wall degrading enzymes (PCWDEs) were found to be differentially expressed in the invasive tissues of both species. Overall the number of upregulated PCWDEs in the invasive mycelium was much higher than the invasive rhizomorphs when compared to their non-invasive counterparts (Fig 5). The saprotrophic *A. cepistipes* had a higher number of DEG/DAP PCWDEs than the pathogenic *A. ostoyae* (Table S6, Fig 5). Among differentially expressed PCWDEs in mycelium vs non-invasive mycelium, upregulated pectinases were most numerous accounting for 17% and 37% of all pectinases in *A. ostoyae* and *A. cepistipes*, respectively. These were followed by cellulases (9%, 27%), hemicellulases (11%, 30%), and expansins (12%, 10%) (Fig 5, see Table S6 for the complete list of differentially expressed PCWDEs). We found most of the lignin-degradation related genes to be downregulated in both species, with none upregulated in *A. ostoyae* and few upregulated ones in *A. cepistipes* (Fig. 5). We found 1 and 8 upregulated LPMOs/GH61 in invasive mycelia of *A. ostoyae* and *A. cepistipes*, respectively, which might act together with the cellobiohydrolases to enhance cellulose degradation [72–77].

**Fig 5.**
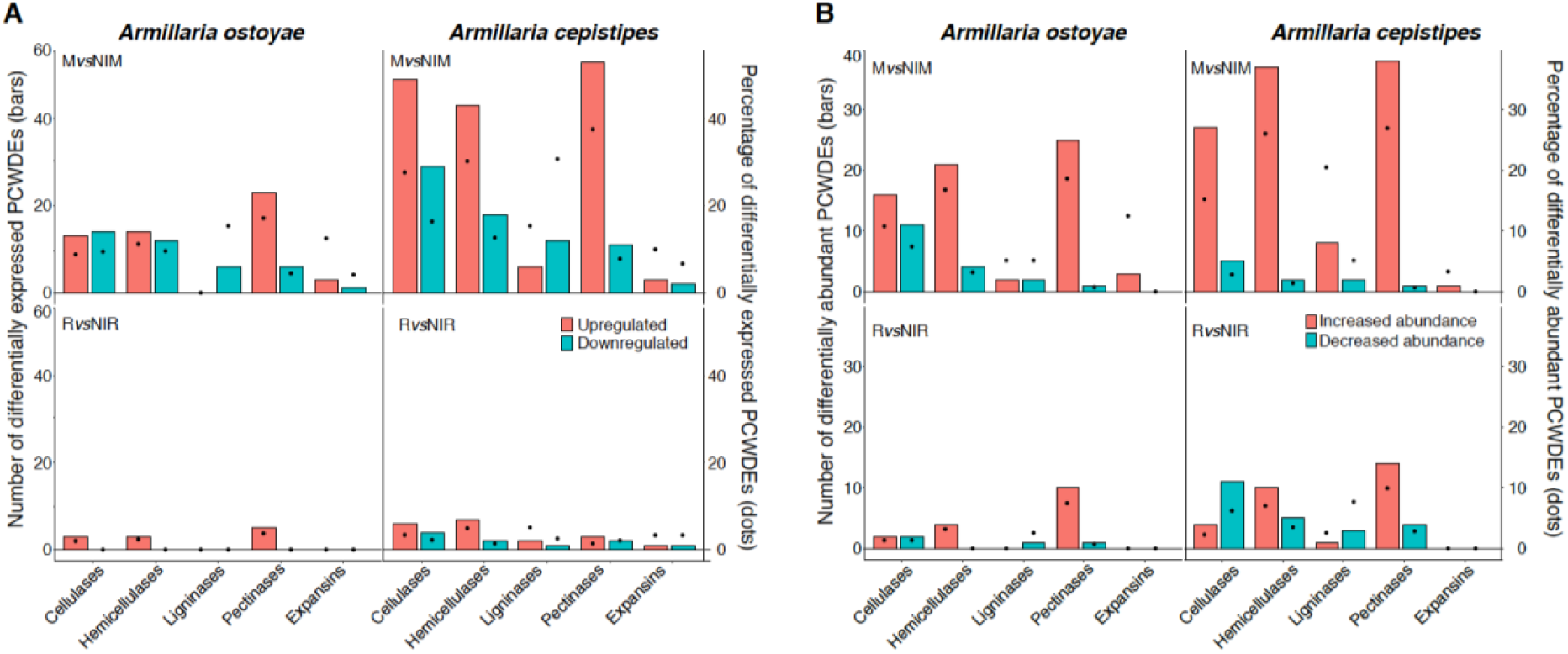
Differentially expressed/abundant plant cell wall degrading enzymes in the two species. A) Barplot showing number, and dots showing the percentage of differentially expressed PCWDEs in the two species in MvsNIM (top) and RvsNIR (bottom) in transcriptomics data. B) Barplot showing the number, and dots showing the percentage of differentially abundant PCWDEs in the two species in MvsNIM (top) and RvsNIR (bottom) in proteomics data. CAZYmes classified on the basis of their substrate in the plant cell wall (Table S6) showing the number of genes upregulated/increased abundance (orange) and downregulated/decrease (blue-green) in the two species.

Our analyses revealed significant expression of both pectinolytic PCWDEs and expansins (Fig 5, Table S6), indicating hallmarks of early-stage wood decay. In previous time-series studies, abundant pectinase expression was found during the early stages of wood-decay, suggesting a requirement of early-stage pectinolytic ‘pretreatment’ [78, 79] for making the plant cell wall structure accessible, followed by a wave of non-pectinolytic GHs expression. In previous studies [78], it was also observed that early stages of wood decay were marked by increased expression of expansins and GH28 pectinases, suggesting both enzymatic and mechanical loosening of the plant cell walls for easier access of the cellulose and hemicellulose components. Our observations of pectinolytic PCWDE and expansin expression are in line with this.

In contrast, we detected few lignin-degrading AAs to be DEG/DAP in the two species, the majority of which were downregulated, suggesting that lignin was not appreciably attacked by *Armillaria* in our experiments. Of the 39 genes that encode lignin degradation related proteins (Table S6) in each of the species, we found no upregulated and about 15% downregulated ligninases in *A. ostoyae*. In *A. cepistipes*, we found 15% upregulated and 30% downregulated ligninases. In the *A. cepistipes*, the proteomic data revealed somewhat more substantial lignin degradation: we found 4 (out of 11 detected) AA1_1 laccases, 3 AA2 peroxidases (out of 4) and 7 (aryl) alcohol dehydrogenases (AA3_2, of 17) increased in invasive relative to non-invasive mycelium. The modest induction and general downregulation (in the transcriptomic data) of ligninolytic genes are remarkable for white-rot fungi, especially in the light of previous studies reporting an early activation of ligninolytic enzymes by white-rot fungi [79, 80] and the enrichment of class II peroxidases and other AAs (e.g. laccases) near the hyphal front. The lack of a ligninolytic burst is, on the other hand, consistent with the underrepresentation of ligninolytic gene families in *Armillaria* compared to other white-rot Agaricales [12]. We note that heat-based lignin breakdown or loosening of the lignocellulose matrix during autoclaving [81–83] may provide a complimentary, although a less likely explanation of our data.

#### Evidence for pectinolysis from galactose binding domain proteins

We found a high number of galactose-binding-like domain superfamily protein (GBDPs) genes upregulated the two species, especially in invasive mycelia. This protein superfamily includes, among others, the rhamnose-binding module family CBM67. Out of 84 and 89 genes containing GBDPs in *A. ostoyae* and *A. cepistipes*, we found 13 and 29 upregulated in invasive vs non-invasive mycelia, respectively. Four of these genes were also among the 20 most highly induced co-orthologous genes in invasive mycelia (see Fig 4). In the proteomics data, *A. ostoyae* had 7 GBDPs (2 CBM67s) and *A. cepistipes* had 12 GBDPs (2 CBM67s) with increased abundance in the mycelium vs non-invasive mycelium. A significant portion (35-50%) of upregulated genes were annotated as CBM67s in the CAZy database [84]. CBM67 are L-rhamnose binding modules, which are reported to be involved in pectin degradation [12, 85]. Apart from CBM67, there were a number of other pectinolytic enzymes (e.g. GH78, PL4) associated with these GBDPs, which were upregulated in the two species (Table S6). The abundance of L-rhamnose binding modules on their own, or in combination with pectinolytic enzyme encoding genes, as well as the dominance of pectinolytic PCWDEs among upregulated CAZymes could suggest a decay strategy focussed on pectin removal for accessing the cellulose and hemicellulose units of the plant cell wall.

#### Iron acquisition genes upregulated in *Armillaria* spp

We observed a number of iron acquisition genes to be upregulated in mycelia vs non-invasive mycelia of both species. Uptake of extracellular ferrous iron (Fe2+) occurs in fungi via a two-part transporter, which consists of an iron permease (Ftr1/FtrA) and an associated multicopper oxidase (Fet3) [86]. In *A. ostoyae* there were 3 Ftr1/Fip1/EfeU iron permease genes (IPR004923) of which 2 were upregulated in the mycelium. Of these 3 iron permeases, two were accompanied by a multicopper oxidase gene (one had an upstream and the other had a downstream MCO to the Ftr1/FtrA iron permease respectively). In *A. cepistipes*, there were 2 iron permease genes, of which one was upregulated in mycelium vs non-invasive mycelium. The genes downstream to these 2 iron permeases were multicopper oxidases, and both were upregulated in mycelium vs non-invasive mycelium. Neither iron permeases nor the Fet3-multicopper oxidases were differentially expressed in rhizomorphs vs non-invasive rhizomorphs. In proteomics, we found 1 Ftr1/FtrA iron permease gene with increased abundance in mycelia of both *A. ostoyae* and *A. cepistipes*. There was 1 downstream multicopper oxidase gene with increased abundance in the mycelium only of *A. cepistipes* and not in *A. ostoyae*. The iron permease system along with iron reductases and MCOs is specifically seen upregulated in brown-rot wood decay [67, 78, 85, 87–89].

#### Diverse cytochrome P450s are differentially expressed in invasive tissues

Cytochrome P450s have diverse functions across the fungal kingdom, including secondary metabolite production, detoxification, aromatic compound degradation, among others [90]. Based on InterPro domains (IPR001128) we found a total of 264 and 307 cytochrome P450 encoding genes in *A. ostoyae* and *A. cepistipes*, respectively. Of these 35 were upregulated and 57 downregulated in invasive mycelia along with 3 upregulated 1 downregulated in rhizomorphs of *A. ostoyae* relative to the non-invasive counterparts of these tissues. In proteomics data for *A. ostoyae*, we found 7 increased and 7 decreased cytP450s in M*vs*NIM, along with 3 increased and no proteins with decreased abundance in RvsNIR. In *A. cepistipes* we found 58 (proteomics: 13) up- and 84 (6) downregulated cytP450s in M*vs*NIM along with 27 (11) up- and 22 (7) downregulated cytP450s in RvsNIR. Most of the cytochrome P450s with increased abundance in invasive mycelia compared to non-invasive were classified as E-class group 1.

Interestingly, among the cytochrome P450 proteins that had decreased abundance in invasive mycelia were co-orthologs of the Psi-producing oxygenase A (*PpoA*) from *Aspergillus nidulans*, which is associated with secondary metabolite biosynthesis (sterigmatocystin), oxylipin biosynthesis and coordination of a/sexual sporulation [91]. Ppo proteins are also implicated in virulence, with *A. fumigatus ppo* mutants displaying hypervirulence, possibly due to the activation of the immune response in mammals [92]. This is observed plant pathogens also, with *Fusarium verticillioides ppo* deletion strain showing higher virulence in assays with maize cobs as well as elevated fumonisin production and lower induction of plant defense-related genes [93]. Therefore, the downregulation of *PpoA* homologs in invasive mycelia of *A. ostoyae* and *A. cepistipes* may result in lower induction of plant defenses, alteration of secondary metabolism, and enhanced virulence.

### Rhizomorphs show an upregulation of transporters as compared to mycelium

*Armillaria* rhizomorphs are putatively involved in the translocation of nutrients [94]. Rhizomorphs are generally believed to serve as migratory organs for exploration of substrates across various distances, however, the studies also indicate rhizomorphs produced by saprotrophic basidiomycetes are also effective in absorbing inorganic nutrients and water from the soil [46, 50–52, 95, 96]. Experiments with *Armillaria mellea* [51, 52] and *Serpula lacrymans* [53, 54, 97] demonstrated the translocation of various nutrients, water, and carbon within the rhizomorphs.

To investigate transporter expression in our wood-decay system, we classified putative transporters in A. ostoyae and A. cepistipes based on conserved domains. We identified 612 and 602 transporters in *A. ostoyae* and *A. cepistipes*, which belonged to 100 InterPro annotations with major facilitator superfamily domain, ABC-transporter like, sugar transporters, amino acid/polyamine transporters, and P-type ATPases being most abundant. Table S9 lists the identified transporters and their expression and abundances in the transcriptomic and proteomic data. Transcriptomics provided dynamics for a much larger number of transporters than proteomics, possibly due to the difficulty of extracting membrane proteins for LC/MS analyses. In the RNA-Seq data, we found 45 and 84 upregulated along with 96 and 105 downregulated transporters in mycelium vs non-invasive mycelium in *A. ostoyae* and *A. cepistipes*, respectively (Fig 6). The majority of transporters upregulated in the mycelium vs non-invasive mycelium belonged to the major facilitator superfamily (MFS) and sugar transporter family. Considerably lower numbers of upregulated transporters were found in rhizomorphs: 2 and 20 upregulated in *A. ostoyae* and *A. cepistipes*, respectively from the same families.

**Fig 6.**
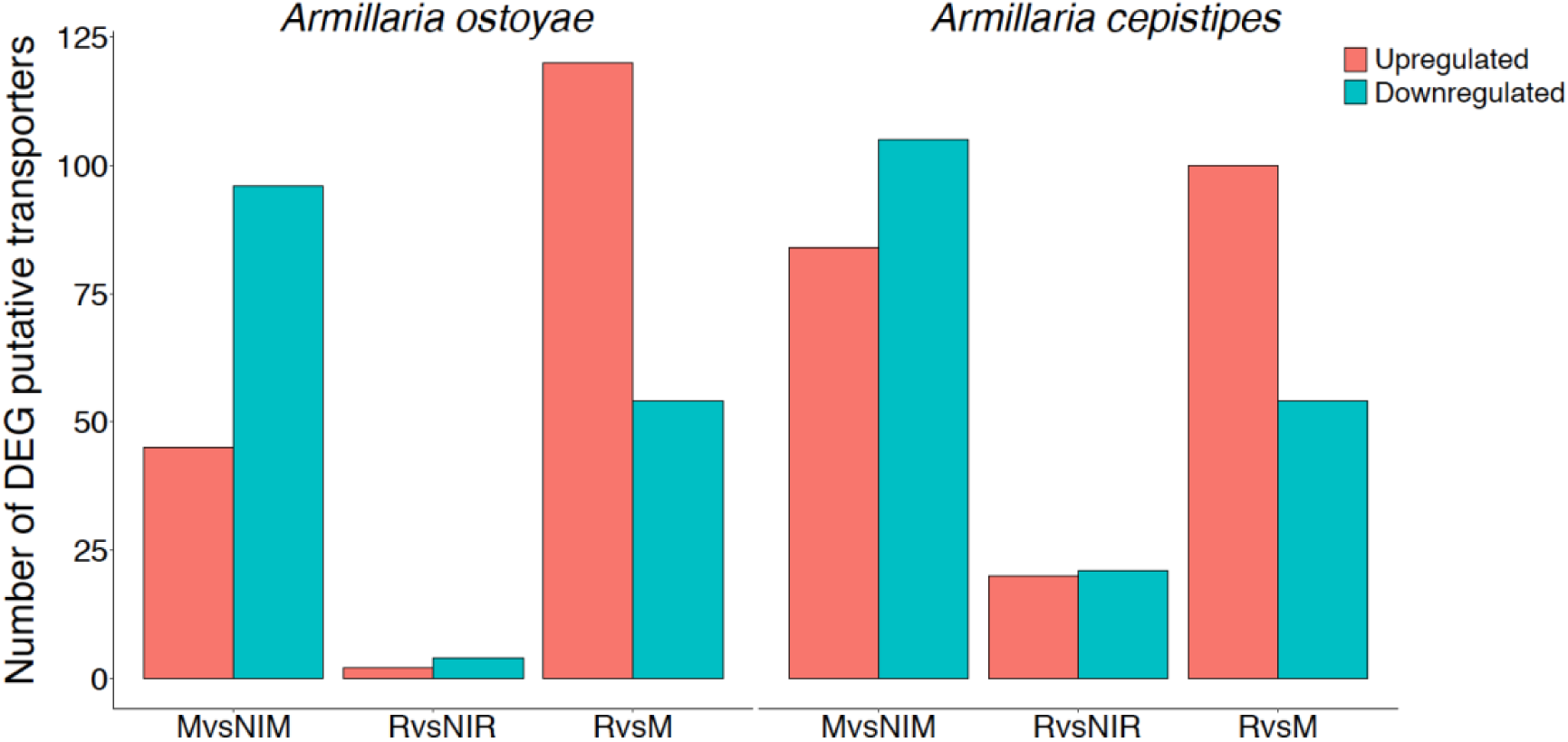
Number of differentially expressed putative transporters in the two species. The number of upregulated (orange) and downregulated (blue-green) genes are shown for M*vs*NIM, R*vs*NIR and R*vs*M comparisons in the two species.

A striking difference in the expression of transporters was found between rhizomorphs vs mycelium (Fig 6), with 120 and 100 upregulated and much fewer downregulated transporters in *A. ostoyae* and *A. cepistipes*, respectively. Compared to the total number of these transporters, the most upregulated transporters in rhizomorphs vs mycelium were the ones possibly involved in sugar transport, such as major facilitator sugar transport-like (IPR005828), sugar/inositol transporter (IPR003663), sugar transporters (IPR005829). Several aquaporin-like proteins, that were reported to be involved in mushroom development [98] and ectomycorrhizal functioning [99], were also upregulated in rhizomorph vs mycelium.

We also compared the upregulated transporters in rhizomorphs vs mycelium to fruiting body development regulated genes of *A. ostoyae* from Sipos et al [12], to identify transporters that are specifically upregulated in rhizomorphs but not in fruiting bodies. We reasoned that such genes might be involved in rhizomorph-specific functions rather than shared multicellularity-related functions between rhizomorphs and fruiting bodies. We found 47 such genes (Table S7), 39% of rhizomorph-upregulated transporters, including several MFS domains and sugar transporters. Collectively, our transporter data suggests rhizomorphs not being involved in active wood-decay rather they might be involved in the transfer of the decomposition intermediates between different parts of the colony.

## Conclusions

In this study, we examined wood-decay patterns by *Armillaria* spp. using transcriptomic and proteomic data on autoclaved spruce roots. Roots were primarily colonized by mycelial tissues, which harbored much more differentially expressed genes and differentially abundant proteins than rhizomorphs, suggesting that, although rhizomorphs can efficiently forage for nutrients, individual hyphae pierce and colonize woody tissues. Of the two species, the saprotroph *A. cepistipes* showed a much stronger molecular response to wood than did the pathogenic *A. ostoyae*. This pattern was evident both in terms of the number of differentially expressed genes/proteins and in gene expression dynamics (e.g. fold change) displayed by the two species. We observed a higher number of upregulated PCWDEs in *A. cepistipes* than in *A. ostoyae* with the latter species showing more down-than upregulation in PCWDEs. We speculate that these observations might reflect a general difference in wood-decay strategies of saprotrophic vs. pathogenic species. Because saprotrophs colonize dead wood, they likely face more intense competition with other microbes than do necrotrophic pathogens, which, after killing the host, are the very first colonizers and thus might face less competition. This might select for more aggressive wood-decay strategy in saprotrophs, which, in an assay like ours may manifest as a stronger induction of PCWDEs. In comparison, pathogens, which can also feed while the host is alive, may not be under a strong pressure to express a large suite of wood-decay enzymes. Similar observations were made in a study comparing gene expression during saprotrophic and parasitic phases in *Heterobasidion irregulare* [32]. It should be noted that these observations might also be influenced by the substrate (though spruce is a natural substrate for both species), the individual properties of the strains, and other factors, so more evidence, and experimental testing are needed to confirm this hypothesis.

*Mycelia of A. ostoyae* and *A. cepistipes* responded similarly to wood, with 713 orthologous genes showing differential expression in both species. These include many plant cell wall degrading enzyme genes, hydrophobins, CBM67s, cytochrome p450s, transcription factors, and iron acquisition-related genes, among others. Pectinolytic PCWDE genes were dominant among CAZymes, followed by expansins, cellulose- and hemicellulose degrading ones, whereas ligninolytic PCWDE genes were mostly downregulated in our assay. The proportionately high number of pectin-related PCWDE-s mirrors comparative genomic observations that revealed enrichment of pectinolytic genes (in particular CBM67s) and expansins, but depletion of ligninolytic ones in *Armillaria* genomes [12]. Of particular interest are CBM67-s, which comprised 4 of the top 20 most induced co-orthologous genes in both species. There was a similar percentage of upregulated pectinolytic genes in the mycelium of *A. cepistipes* (ca. 38%) as in other studies examining early-phase decay (48% in *Pycnoporus coccineus* [100] *and 45% in Postia placenta* [78]*), whereas the percentage was lower in A. ostoyae* (17%). Ligninolytic genes, on the other hand, were underrepresented among DEGs/DAPs, consistent with their general underrepresentation in *Armillaria* genomes. This aligns well with previous reports of the limited lignin-degrading capacity of *Armillaria* spp. [38, 39, 101].

Several aspects of the gene expression patterns in our assays are unusual for white-rot fungi. These include the lack of an early ligninolytic gene expression burst as is typical for white-rot fungi and the high expression of some genes (e.g. iron uptake systems, oxoglutarate/iron-dependent dioxygenases) that have been reported from brown rot fungi [67, 78, 85, 87–89]. Previous studies questioned the typical white-rot nature of *Armillaria* [36, 38, 39, 101]. *It was observed, through chemical analysis, that Armillaria mellea* attacked celluloses in the early stages of decay, but not lignin [39, 101] and thus inferring that *Armillaria* performs a variety of white rot in which lignin degradation is not the primary concern. Almasi et al [102] also showed that, based on lignin-degrading genes, *Armillaria* spp. are intermediate between brown-rot and white-rot fungi, along with other species (*Schizophyllum commune, Auriculariopsis ampla, Cylindrobasidium torrendi*), that have been recalcitrant to this dichotomous classification [36, 43], as well as ectomycorrhizal fungi with reduced ligninolytic repertoires. Of these, *C. torrendii* is a close relative of *Armillaria* in the Physalacriaceae. Floudas et al showed that the reduced ligninolytic gene repertoire of *C. torrendii* is a result of gene loss compared to its white-rot ancestors and that the decay caused by this species resembles soft rot. While soft rot in the strict sense is characteristic of the Ascomycota, several Basidiomycetes have been associated with Type II (e.g. *C. torrendii* [36]*) or Type I (e*.*g. Armillaria* spp. [38]) soft rot. These species are characterized by a complete set of PCWDEs for degrading (hemi)cellulose, but a depletion of ligninolytic ones. More generally, a selective deployment of ligninolytic activity has been reported in a number of species. *Mucidula mucida* (also Physalacriaceae, as *Oudemansiella mucida)* and *Meripilus giganteus* (Polyporales) caused either soft or white rot, depending on decay stage and substrate (host species and cell type) [103, 104].

Wood decay resembling soft rot has been reported also for several early-diverging Agaricomycetes (e.g. in non-mycorrhizal Cantharellales), which predate the origin of ligninolytic class II peroxidases [17, 43, 44]. We hypothesized that early diverging Agaricomycetes and more derived species that lost some of their ligninolytic but not their cellulolytic gene repertoires (e.g. *Jaapia, Schizophyllum*), reverted to a plesiomorphic soft-rot like decay chemistry, which is primarily dominated by cellulolytic and pectinolytic functions [44]. It appears that this is a particular characteristic of the Physalacriaceae (e.g. *Armillaria, Mucidula, Cylindrobasidium*), which lost some ligninolytic genes compared to their white-rot ancestors but evolved mechanisms for selectively deploying the remaining ligninolytic genes based on environmental factors. We think that the combination of (i) a widely conserved plesiomorphic soft rot-like wood decay strategy and (ii) the ability to degrade lignin in white rot, enables Agaricomycetes to toggle between soft- and white rot either by gene loss or by gene expression regulation. Thus, it is possible that temporal or substrate-dependent regulation of the activation of ligninolysis can separate soft- and white-rot behaviors of some species, adding further complexity to the range of decay modes of Basidiomycota.

## Methods

### Wood colonization assay and RNA-Extraction

Cultures of *Armillaria ostoyae* C18 and *Armillaria cepistipes* B2 [12] were inoculated on Malt extract agar (MEA) and incubated at 25°C in dark for a week. 4-5 cm long, autoclave sterilized spruce roots were introduced to the week-old cultures of *A. ostoyae* and *A. cepistipes* and were again kept at 25°C in the dark for 2-3 weeks until they were colonized by the fungi (Fig 1B). After colonization, roots were dissected to collect mycelium from below the outer bark layer and rhizomorphs emerging out of the colonized wood (Fig 1B). Cultures of *Armillaria* without the addition of sterilized spruce roots were used as non-invasive controls. Total RNA was extracted from the four tissue types in three biological replicates, using the Quick-RNA Miniprep kit (Zymo Research, Irvine, CA, USA), following the manufacturer’s protocol.

### RNA-Seq library generation and sequencing

RNA-Seq analyses were carried out by using Ribo-Zero rRNA removal kit (Yeast) to deplete rRNA from total RNA. Subsequently, samples were processed with Illumina TruSeq V2 library preparation protocol. Libraries were sequenced on an Illumina NextSeq 500 machine yielding 2×150 nt reads.

### Protein extraction

For protein extraction, tissues were snap-frozen in liquid N_2_ and bead beaten periodically (30 Hz, 2 min), with snap freezing between cycles. Lysis buffer (6M Guanidine-HCl, 0.1 M Tris-HCl, 50 mM DTT pH 8.6) was added to crushed fungal tissue and bead-beating was repeated. Samples were further disrupted using sonication (MS72 probe, 3 x 10 sec), with cooling on ice between sonications. Samples were clarified by centrifugation and supernatants passed through 3 kDa cut-off filters (Millipore) to concentrate and perform buffer exchange into PBS. Protein samples were precipitated with TCA (final 15 % w/v) and pellets were washed with ice-cold acetone. Protein pellets were resuspended in UT buffer (6 M Urea, 2M Thiourea, 0.1 M Tris-HCl pH 8) and concentrations normalized following Bradford protein assay. Samples were digested according to Moloney et al [105] and ZipTips (Millipore) were used for sample clean-up. Peptide samples were analyzed using the high mass accuracy Q-Exactive mass spectrometer coupled to a Dionex Ultimate 3000 nanoLC with an EasySpray PepMap C18 column (50 cm × 75 µm). Peptide mixtures were separated as described in Collins et al [106] and resultant data were analyzed using MaxQuant (v 1.5.3.30) [107] with the label-free quantitation (LFQ) algorithms and searching against the protein database (filtered models) in JGI MycoCosm [12].

### Bioinformatic analyses of RNA-Seq data

Paired-end Illumina (HiSeq, NextSeq) reads were quality trimmed using the CLC Genomics Workbench tool version 11.0 (CLC Bio/Qiagen) removing ambiguous nucleotides as well as any low quality read end parts. The quality cutoff value (error probability) was set to 0.05, corresponding to a Phred score of 13. Trimmed reads containing at least 40 bases were mapped using the RNA-Seq Analysis 2.16 package in CLC requiring at least 80% sequence identity over at least 80% of the read lengths; strand specificity was omitted. List of reference sequences is provided in Table S1 along with the mapping statistics for both species. Reads with less than 30 equally scoring mapping positions were mapped to all possible locations while reads with more than 30 potential mapping positions were considered as uninformative repeat reads and were removed from the analysis. “Total counts” RNA-Seq count data was imported from CLC into R version 3.0.2. Genes were filtered based on their expression levels keeping only those features that were detected by at least five mapped reads in at least 25% of the samples included in the study. Subsequently, “calcNormFactors” from “edgeR” version 3.4.2 [108] was used to perform data scaling based on the”trimmed mean of M-values” (TMM) method [109]. Log transformation was carried out by the “voom” function of the “limma” package version 3.18.13 [110]. Linear modeling, empirical Bayes moderation as well as the calculation of differentially expressed genes were carried out using “limma”. Genes showing an at least two-fold gene expression change with an FDR value below 0.05 were considered as significant. Multidimensional scaling (“plotMDS” function in edgeR) was applied to visually summarize gene expression profiles. In addition, unsupervised cluster analysis with Euclidean distance calculation and complete-linkage clustering was carried out on the normalized data using “heatmap.2” function from R package “gplots”.

### Data availability

RNA-Seq data was deposited in the NCBI’s Gene Expression Omnibus (GEO) Archive at www.ncbi.nlm.nih.gov/geo (accession no. GSE149732).

### Analyses of proteomic data

Proteomic results were organized and statistical analyses were performed using Perseus (v 1.5.4.0) [111]. Qualitative and quantitative analyses were performed to determine the relative changes in protein abundance in each of the sample types. Quantitative analysis was performed using a student’s t-test with a p-value cut-off of 0.05 and log_2_(fold change; FC) ≥ 1. Qualitative analysis revealed proteins that were detected in 2/3 replicates for a sample type and undetectable in all replicates of the comparator group. A theoretical minimum fold change was determined for qualitative results based on a calculated minimum detectable protein intensity (mean + 2 standard deviations of lowest detectable protein intensity for each replicate in the experiment) [105]. Based on this theoretical minimum fold change, some qualitative results were excluded due to intensity values approaching the minimum detectable levels. Qualitative and quantitative results were combined and a total number of differentially abundant proteins (DAPs) are summarised in Table S3 and Fig 1E.

### Clustering analysis and functional annotation

Predicted protein sequences from JGI MycoCosm were used to find the orthologs in *A. ostoyae* and *A. cepistipes* by OrthoFinder v2.3.1 [62] (default parameters). Single-copy orthologs for the two species (Table S5) were further analyzed for comparing the expression patterns in the two species. Functional annotation was carried out based on Interpro domains using InterProScan v5.24-63.0 [112].

Enriched GO terms for different comparisons were predicted by the topGO package [113], using the weight01 algorithm and Fisher testing. Terms with p-values less than 0.05 were considered significant and were plotted with respect to their gene ratios in (Fig 2, Fig S4), where gene ratio is the number of a particular GO term in a specific comparison type to the total number of that term found in the gene list for that organism (Table S4).

The CAZyme copy numbers in *A. ostoyae* and *A. cepistipes* were collected from the JGI mycocosm annotations, which were based on the CAZy annotation pipeline [84]. We separated the CAZy families based on their substrate-specific plant cell wall degradation abilities (Table S6) and analyzed the copy numbers of differentially expressed genes and differentially abundant proteins for cellulases, hemicellulases, pectinases, expansins and ligninases in the M*vs*NIM and R*vs*NIR comparisons.

Transporters were identified by using DeepLoc [114] to select plasma membrane-localized proteins from the proteomes of both species. Plasma membrane-localized proteins with more than 1 transmembrane domains were used to obtain a list of non-redundant InterPro domains, which were manually checked for functional roles in transport.

## Acknowledgments

The authors acknowledge support by the Hungarian National Research, Development and Innovation Office (contract No. GINOP-2.3.2-15-2016-00052), the ‘Momentum’ program of the Hungarian Academy of Sciences (contract No. LP2019-13/2019 to L.G.N.) and the European Research Council (grant no. 758161 to L.G.N.). Proteomics instrumentation was funded by a competitive Science Foundation Ireland (SFI) Infrastructure award (12/RI/2346 [3])

## Supplementary Figures

**Fig S1.**
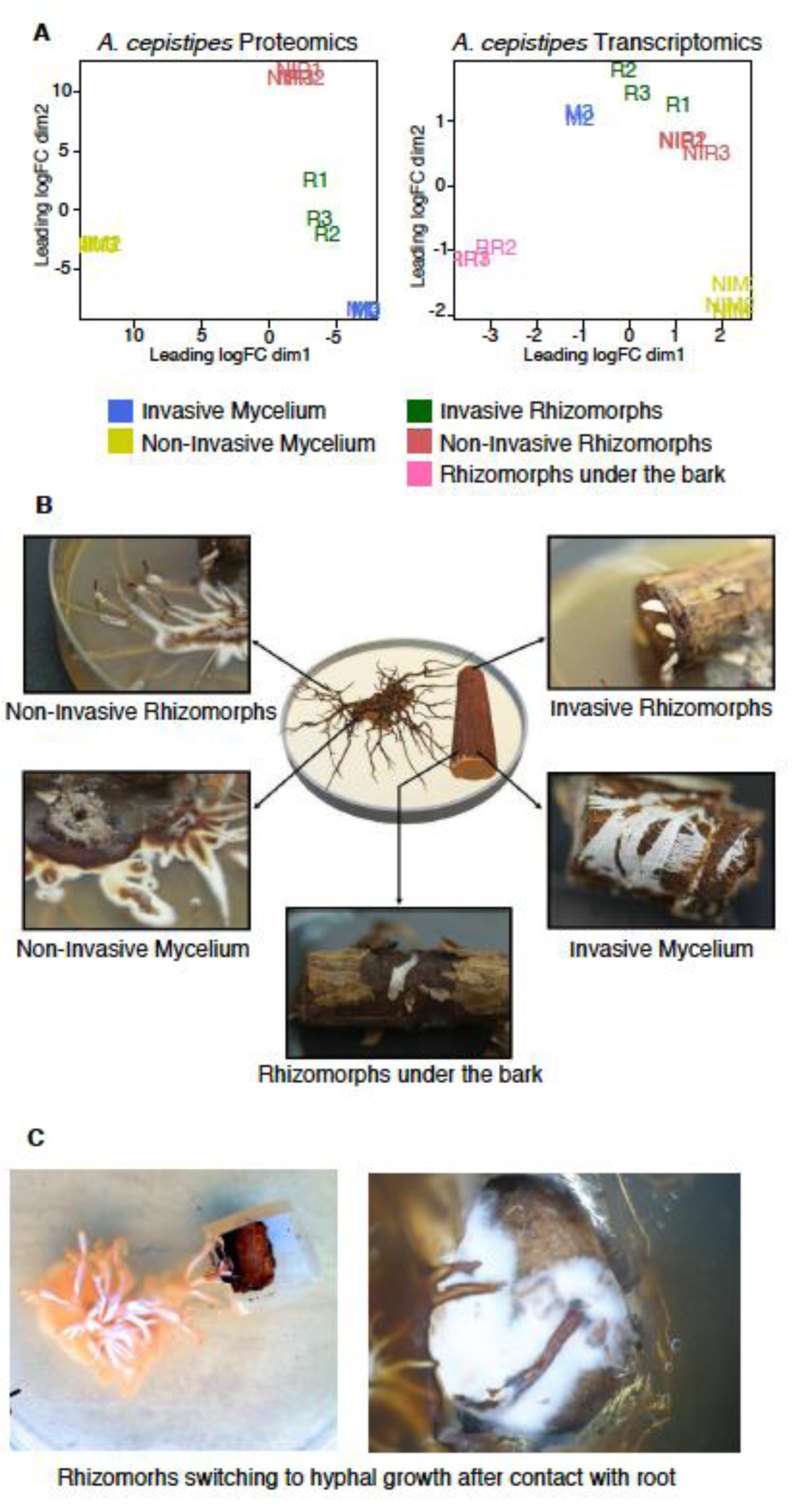
Experimental setup. A) Multidimensional scaling of three biological replicates from each of the tissue types in *A. cepistipes* for proteomics (left) and transcriptomics (right). B) The four tissue types sampled for transcriptomics and proteomics analysis viz. invasive mycelium (growing beneath the outer layer of the root), invasive rhizomorphs (emerging out of the roots), non-invasive mycelium and non-invasive rhizomorphs (growing in absence of root), along with additional RR (rhizomorphs growing beneath the outer layer of the root) in *A. cepistipes*. C) Pictures showing rhizomorphs differentiating into hyphae in contact with the spruce root.

**Fig S2.**
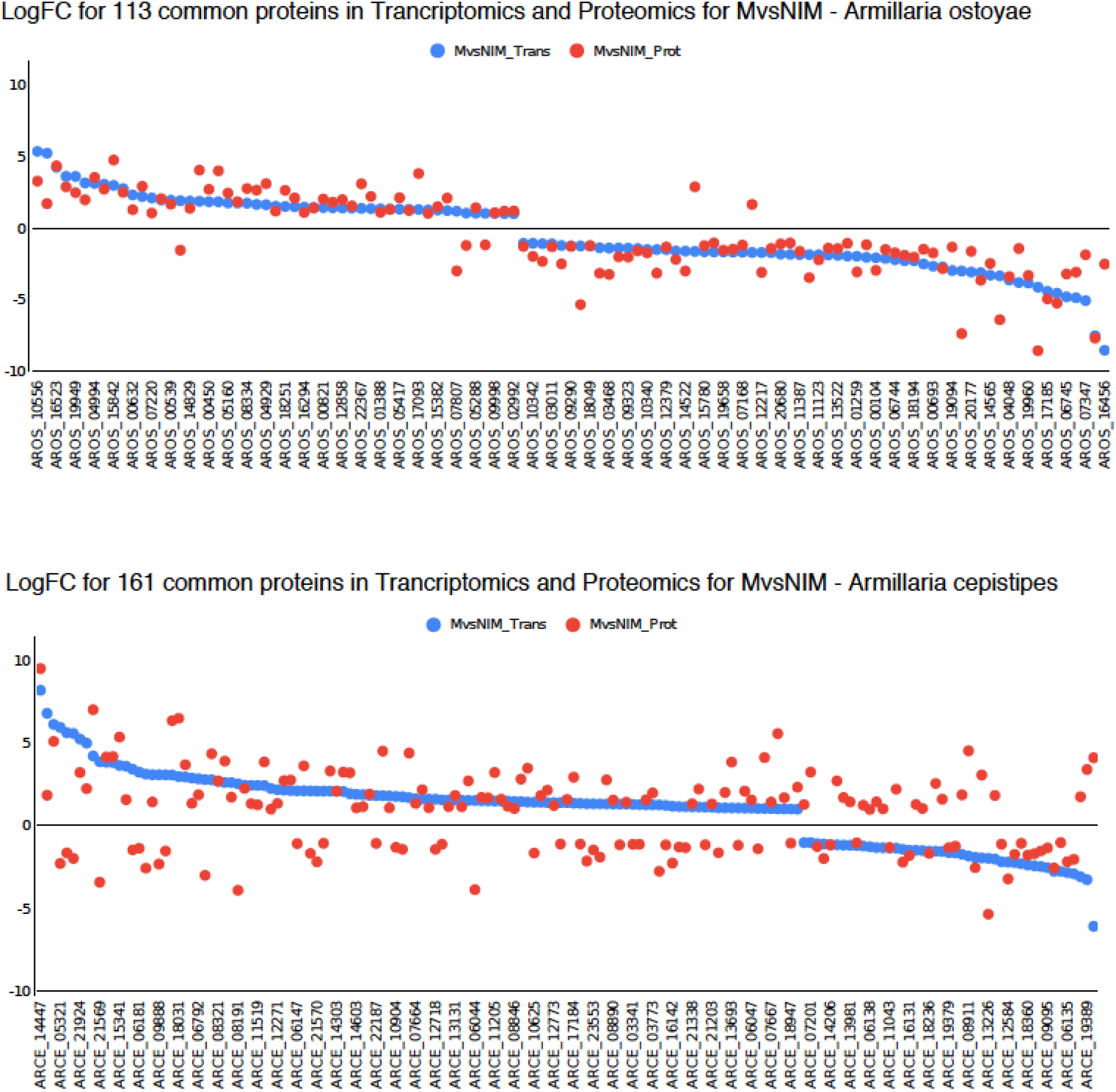
Log fold changes in MvsNIM of genes identified in both transcriptomics and proteomics data in *A. ostoyae* (top) and *A. cepitsipes* (bottom). The logFC for transcripts (blue) are arranged from increased to decreasing order, overlaid with logFC from proteomics (red) shows a limited correlation between the two omics approaches.

**Fig S3.**
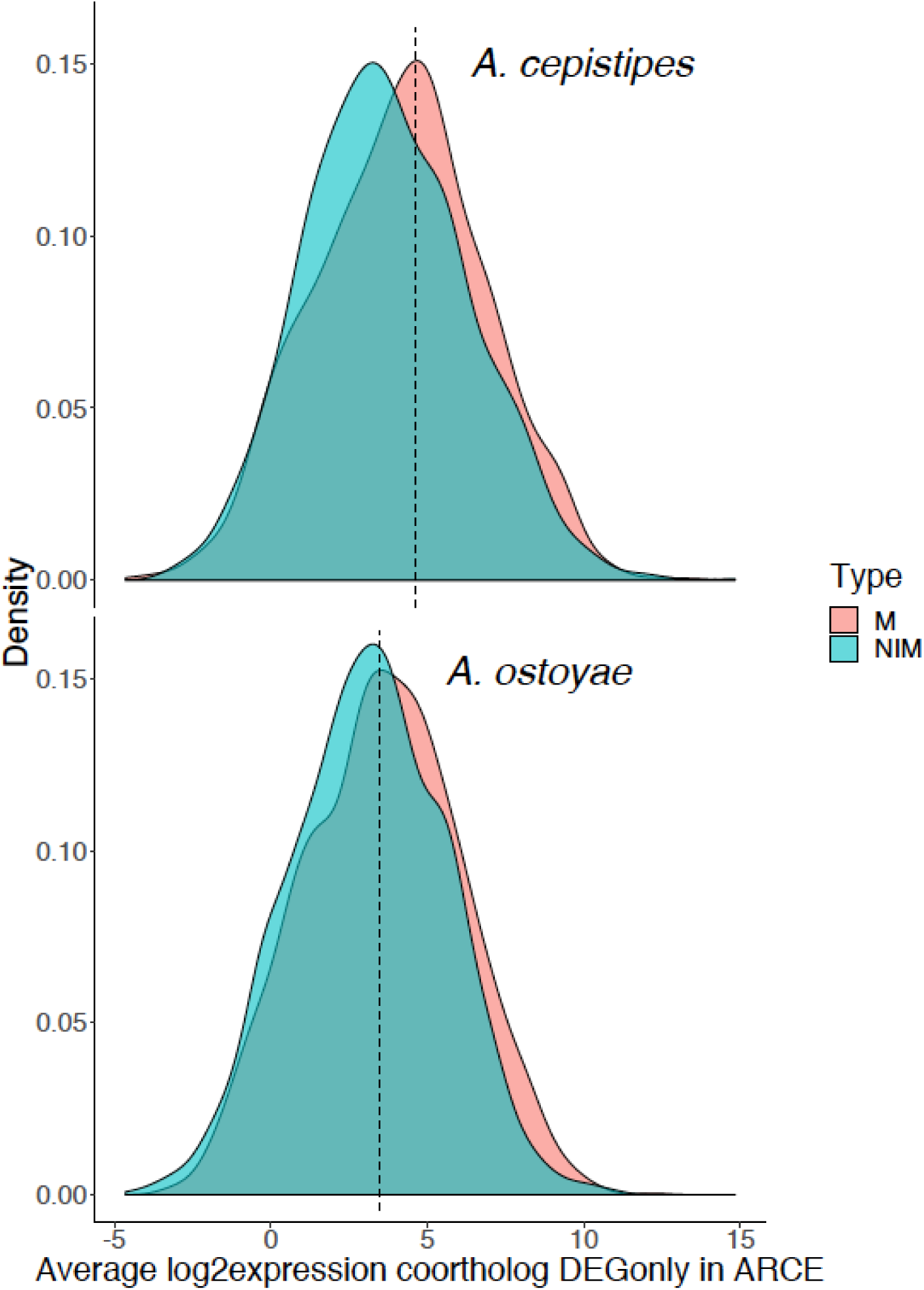
Distribution of raw expression values in both species for co-orthologs differentially expressed in MvsNIM of *A. cepistipes*. Baseline expression of genes in non-invasive mycelia of *A. ostoyae* was not higher than that in A. cepistipes, indicating a stronger response of *A. cepistipes*.

**Fig S4.**
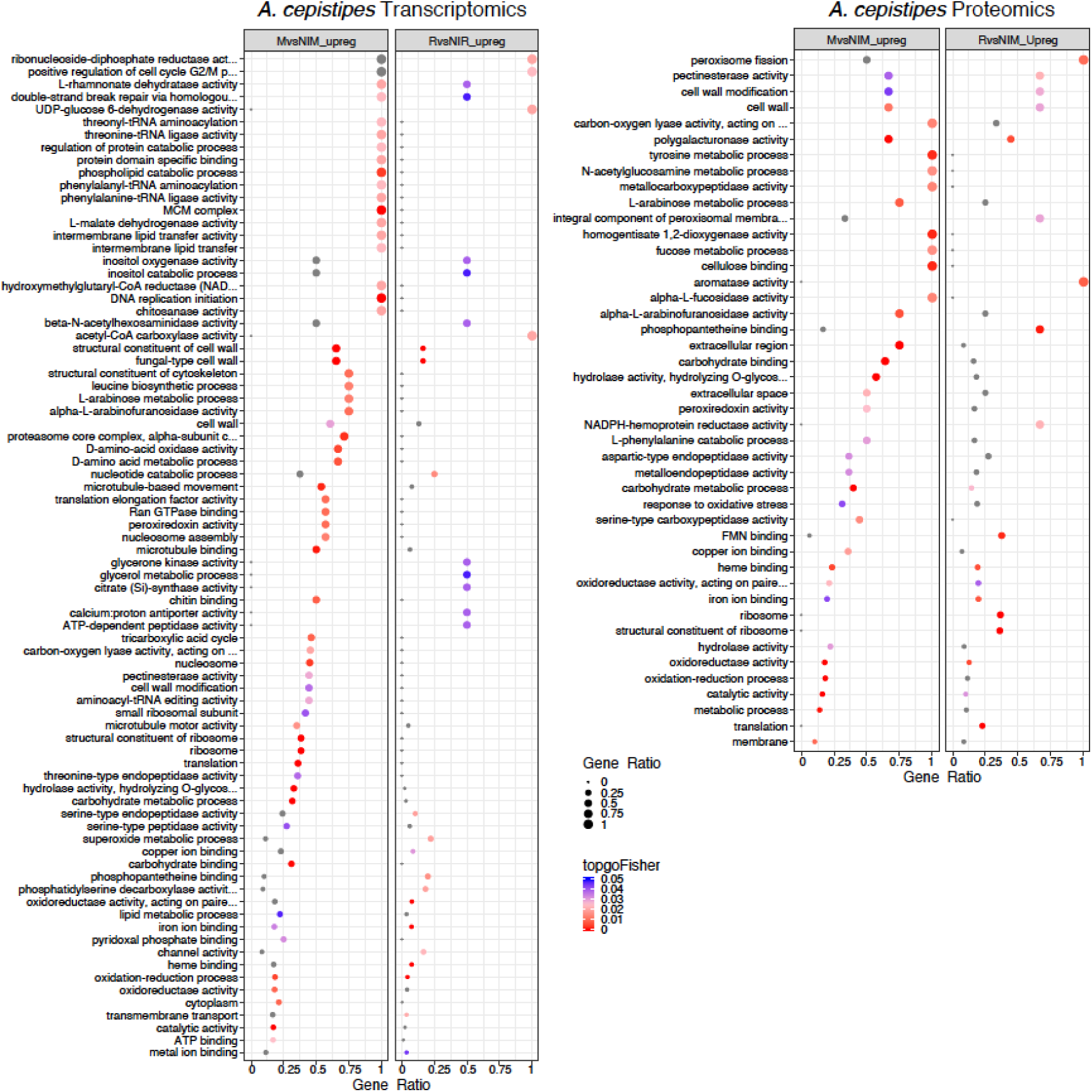
Enriched GO terms in M*vs*NIM and R*vs*NIR of *A. cepistipes* for transcriptomics (left) and proteomics (right). The ratio of number of a particular GO term in a specific comparison (mycelium vs non-invasive mycelium or in rhizomorphs vs non-invasive rhizomorphs) to the total number of that GO term for a species was used to plot gene ratios for enriched GO terms (p<0.05, Fisher’s exact test). The size of the dot is directly proportional to gene ratio, and the color of the dots corresponds to p-values. Grey dots represent GO terms, enriched in only one of the comparisons *i*.*e* either mycelium vs non-invasive mycelium or rhizomorphs vs non-invasive rhizomorphs.

**Fig S5.**
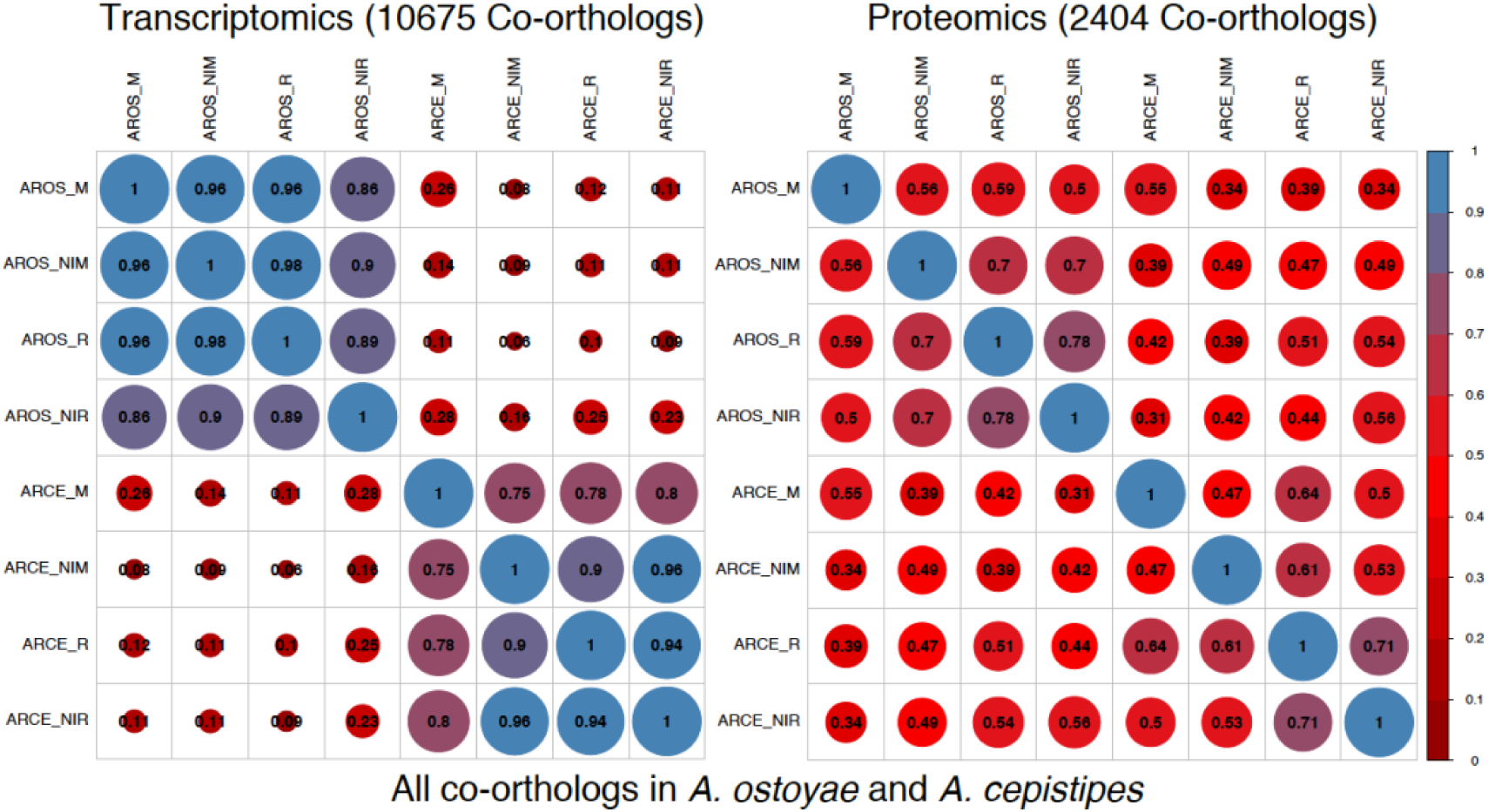
Correlogram for all co-orthologs in the two species, showing correlation between samples across the two species. Blue represents a higher correlation and red represents lower. The size of the circle is directly proportional to a higher correlation. Pairwise mean Pearson correlation coefficients are indicated as numbers in the circles.

## Supplementary Tables

Table S1. RNA-Seq mapping statistics for *A. ostoyae* and *A. cepistipes*

Table S2. Differentially expressed genes (RNA-Seq) in *A. ostoyae* and *A. cepistipes*

Table S3. Differentially abundant proteins (Proteomics) in *A. ostoyae* and *A. cepistipes*

Table S4. List of enriched GO terms in *A. ostoyae* and *A. cepistipes* in the transcriptomics and proteomics analyses

Table S5. Single copy co-orthologs in both species, with common and species-specific DEGs/DAPs in *A. ostoyae* and *A. cepistipes*

Table S6. Carbohydrate-active enzymes (CAZymes) and plant cell wall degrading enzymes (PCWDEs) identified in transcriptomics and proteomics analyses of *A. ostoyae* and *A. cepistipes*

Table S7. Putative transporters identified in transcriptomics and proteomics analyses of *A. ostoyae* and *A. cepistipes*

